# Modifications to gene body methylation does not alter gene expression plasticity in a reef-building coral

**DOI:** 10.1101/2023.09.25.559388

**Authors:** Evelyn Abbott, Coral Loockerman, Mikhail Matz

## Abstract

As coral reefs continue to decline due to climate change, the role of coral epigenetics (specifically, gene body methylation, GBM) in coral acclimatization warrants investigation. The evidence is currently conflicting. In diverse animal phyla the baseline GBM level is associated with gene function: continuously expressed “housekeeping” genes are typically highly methylated, while inducible context-dependent genes have low or no methylation at all. Some authors report association between GBM and the environment, and interpret this observation as evidence of GBM’s role in acclimatization. Yet, others argue that correlation between GBM change and gene expression change is typically absent or negligible. Here, we used the reef-building coral, *Acropora millepora,* to test whether environmentally driven changes in GBM are associated with a gene’s ability to respond to environmental changes (plasticity) rather than expression level. We analyzed two cases of modified gene expression plasticity observed in a three-week-long heat acclimatization experiment. The first one was a group of heat-induced genes that failed to revert their expression after the coral was translocated back to the control tank. The second case involved genes that changed the magnitude of their response to the daily temperature fluctuations over the course of the experiment. In both cases we found negligible or no association with GBM change. We conclude that although both gene expression plasticity and GBM can change during acclimatization, there is no direct association between the two. This adds to the increasing volume of evidence that the function of GBM in invertebrates is unrelated to acclimatization on physiological timescales.

## Introduction

Ocean warming due to anthropogenic climate change has caused increasingly consistent, severe coral bleaching events, leading to widespread reef decline. There has been a growing interest in epigenetics and how it may facilitate adaptation in a rapidly changing environment. However, there is conflicting evidence that modifications to the epigenome serve an adaptive function in cnidarians.

DNA methylation is a common epigenetic modification among eukaryotes which may have originated as a mechanism for genomic defense against transposable elements (Chandler et al 1986; Hackett et al 2013). In animals, it is highly heritable (Bird 2002) and maintained during cell division by DNA methyltransferases, which are homologous among vertebrates and invertebrates (Lyko 2017). DNA methylation in vertebrates occurs throughout the genome, including promoters, enhancers, and gene bodies. Promoter methylation serves a variety of functions, including gene silencing, X-inactivation, gene imprinting, and cell differentiation during development (Moore et al, 2013; Smith & Meissner, 2013). Methylation of gene bodies, on the other hand, is linked to elevated gene expression (Jones 2012).

In invertebrates, DNA methylation levels vary greatly between taxa and occurs almost exclusively in gene bodies (Suzuki et al., 2007). Gene body methylation (GBM) level has a bimodal distribution across the genome and is correlated with how consistently a gene is expressed. Genes with high GBM are constitutively expressed, while genes with low GBM are dynamically regulated and tend to be expressed at lower levels (Jones et al., 2012; Suzuki et al., 2007; Zemach et al., 2007). This raises the question of whether the environment can change GBM and if this change can alter gene expression.

Previous studies in many invertebrates have demonstrated that GBM may be subject to change, including species of bees, ants, moths, and crayfish (Gatzmann et al., 2018; Harris et al., 2019; de Mendoza et al., 2020). This has also been demonstrated in cnidarians, such as anemones and corals (Dixon et al., 2022). However, the potential function of these changes remains controversial. One study which investigated ocean acidification acclimatization in the coral, *Stylophora pistillata,* found that higher intensity stress was associated with increased GBM and decreased transcriptional noise (Liew et al 2018). Another study found that in the coral, *Platygyra daedalea*, GBM patterns were associated with specific life cycle stages and that GBM changes were associated with offspring survival (Liew et al 2020).

On the contrary, other studies have come to different conclusions. A study of the crayfish, *Procambarus virginalis*, found no variation in GBM across tissue types (Gatzmann et al., 2018), despite profound differences in gene expression. Another study conducted on European honey bees found that while GBM varies during development, this variability has no association with gene expression change (Harris et al., 2019). In other insects, DNMT gene knock out lineages had no transcriptional differences compared to their wild type counterparts (de Mendoza et al., 2020). Furthermore, a meta-analysis spanning eight invertebrate taxa, including two cnidarians (the coral, *Stylophora pistillata*, and the anemone, *Exaiptasia pallida*), found no biologically-relevant correlation between GBM change and gene expression (Dixon et al., 2022).

Here, we hypothesized that environmentally-driven GBM changes may not affect the level of gene expression, but rather the plasticity of gene expression, i.e., the ability of a gene to change its expression in response to environmental perturbation. Specifically, we envisioned that modification of GBM during acclimatization might prevent genes that changed their expression from reverting back to their original state. Our hypothesis was that acclimatization-related GBM change would influence gene expression plasticity, which could be a mechanism by which the organism could “learn” to maintain certain gene expression levels despite short-term environmental fluctuations. To test this hypothesis, we conducted a two-week heat acclimatization experiment with the reef-building coral, *Acropora millepora*, to induce GBM change. We then moved a subset of the heated corals to the control condition for a day to test whether genes with changed GBM remained in the “heated” state. Because high GBM is observed in genes with more stable expression, we expected that increasing methylation would reduce gene expression plasticity.

## Materials and methods

### Sample sourcing and experimental design

The experiment was conducted at Orpheus Island Research Station in November 2018 under GBRMPA permit G18/41245.1. Three *Acropora millepora* colonies were collected by SCUBA at Northeast Orpheus and placed in raceways with unfiltered seawater. The corals were then fragmented into 5-7 cm fragments containing 2-3 tips (“twigs”) and returned to the raceway. For the experiment, twigs were hung from a fishing line in two 70 L containers with a slow flow- through of filtered seawater and aeration to create water movement. The experiment was conducted on shaded tables outside to allow for daily temperature fluctuations. At the start of the experiment, the heater for the heat container was set to 29.5 °C. Due to unexpected warm weather, on day 7 the heater was raised to 31 °C to create a greater contrast with the control temperature. The precise temperatures of the heat and control containers throughout the experiment are shown in Figure S4.

On day four, 12 samples from the heat group were switched to the control container and allowed to acclimate for 24 hours. At 8:00am on day five, 6 samples were taken from each of the heat, control, and switch groups, followed by another 6 samples from each group at 4:00pm. This process was repeated on day 13 and another 12 samples from all groups on day 14, six from each at 8:00am and 4:00pm. Samples were immediately fixed in 100% ethanol and stored at -80 °C.

**Figure 1.**
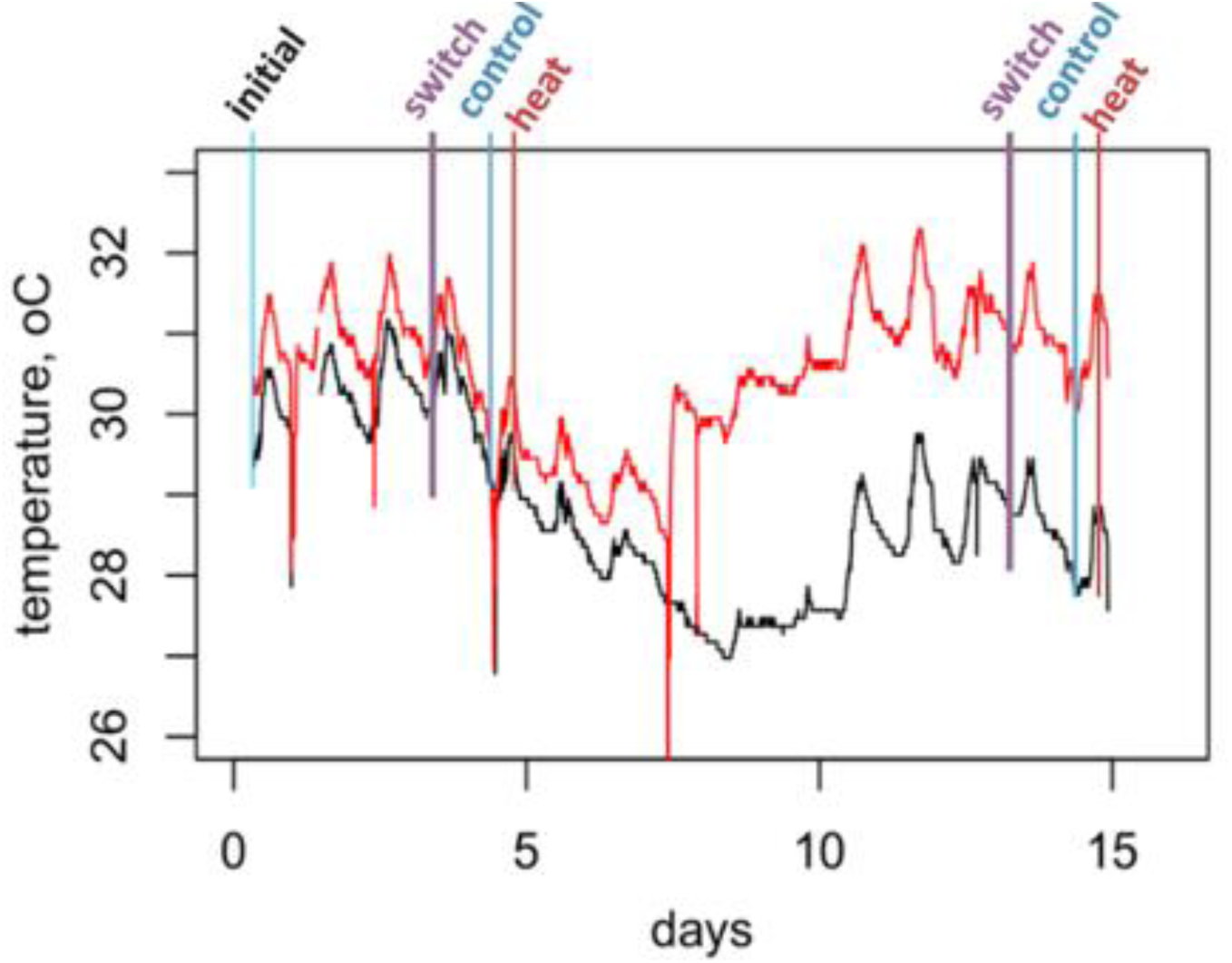
Temperature of control (blue), switch (purple) and heat (red) samples at the first and second experimental timepoints. The deep downward spikes are artifacts of briefly taking loggers out of the tanks for checking.

### Nucleic acid extraction and library preparation

RNA was extracted using an RNAqueous total RNA isolation kit. Samples were submerged in a lysis buffer and pulverized using 150-212uM glass beads and Biospec Mini-Beadbeater for 20 seconds. The RNA was then extracted following the kit’s protocol. DNA was isolated from the same lysate using a phenol:chloroform:isoamyl alcohol extraction, followed by purification with a Zymo cleanup and concentrator kit. Details of the DNA extraction protocol can be found at https://github.com/evelynabbott/DNA_methylation_plasticity. TagSeq libraries for gene expression analysis (Meyer et al., 2011) were prepared according to the protocol hosted on the TagSeq github page, https://github.com/z0on/tag-based_RNAseq. Gene body methylation (GBM) was analyzed using mdRAD protocol (Dixon & Matz, 2020). Detailed protocols can be found in the supplementary materials. BluePippin (Sage Science) was used to size select for fragments 350-550 bp in length, and all samples were sequenced at a depth of 2-3 million reads per sample. Sequencing was conducted at the Genome Sequencing and Analysis Facility at the University of Texas at Austin.

### Data processing

Details of the data processing pipeline and scripts used can be found at https://github.com/evelynabbott/DNA_methylation_plasticity. Adapters were trimmed in single-end mode using cutadapt (Martin, 2011), with a minimum length of 20 bp and a PHRED quality cutoff of 20. FASTQC (Andrews, 2010) was used to check the quality of a subset of 10,000 reads before and after trimming. Reads were mapped to *Acropora millepora* reference genome (Fuller, 2020) using bowtie2 with the --local option, and PCR duplicates were identified using MarkDuplicates from the Picard toolkit (Broad Institute, 2019). Sam files were sorted and converted to bam files using Samtools (Li et al., 2009). mdRAD reads mapping to annotated gene boundaries were counted using FeatureCounts (Liao et al., 2014), and BEDTools Multicov was used to determine fold coverage in different region types (such as transcriptional start sites, promoter boundaries, gene boundaries, etc.). The number of mdRAD counts per sample ranged from 569,974 to 1,321,009, and the number of TagSeq counts per sample ranged from 235,588 to 825,559. The results from BEDTools Multicov were used for mdRAD statistical analyses. Fragments Per Kilobase of transcript per Million mapped reads (FPKM) averaged across all samples was used to calculate gene methylation level.

### Differential gene expression and methylation analysis

To investigate the relationship between gene expression and GBM change due to treatment, experimental timepoint, and time of day, TagSeq and mdRAD counts were imported into DESeq2 (Love et al., 2014) to generate normalized variance-stabilized data. To compute log_2_ fold differences, treatment (control, heat, or switch), experimental time point (1 or 2), and time of day (morning or afternoon) were used as predictive variables.

This DESeq2 formula was also used to analyze the relationship between daily plasticity and GBM change. To compute daily plasticity at time 1 and time 2, log_2_ fold expression changes due to the time of day were computed separately for each time point. DESeq2 was then repeated with all the samples combined, but with the addition of the interaction between time of day and experimental time point. The log_2_ fold changes computed from the interaction term represent the change in daily plasticity over time.

Contour lines representing the effect of GBM log_2_ fold changes and GBM basemean (the average of normalized count values, dividing by size factors, across all samples) on gene expression were calculated using the function *ordisurf()* from the R package *vegan* (Oksanen et al., 2020). *ordisurf()* fits ordination plots with contour lines representing the effect of a given variable using generalized additive modeling.

### Weighted gene co-expression network analysis (WGCNA)

To identify modules of genes co-regulated due to treatment, we used WGCNA (Langfelder & Horvath, 2008). The input for this analysis was a matrix of variance stabilized counts generated by DESeq2 with the effect of genotype removed using the R package limma. Soft threshold was set at 12 and the cut height was set at 0.4, generating 9 modules (Fig. 2).

**Figure 2.**
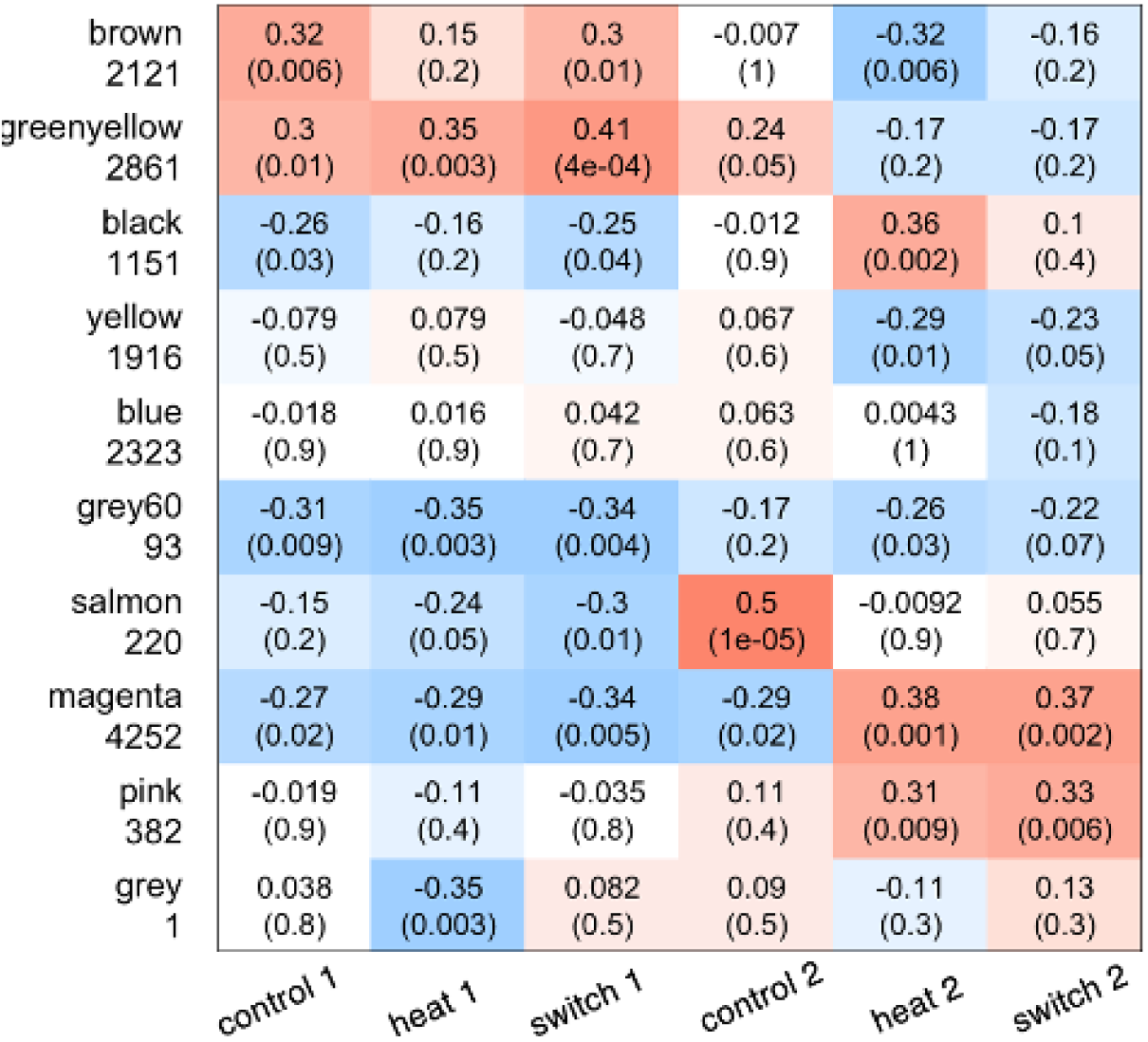
Expression of WGCNA modules in all treatment groups (control, heat, or switch) and time points (1 or 2). Red indicates upregulation and blue indicates downregulation. Row names correspond to module names and the number of genes in the module. The numbers in each cell represent the significance of module expression in each treatment group (top is R, bottom is p value).

One module, “magenta” (4342 genes), was determined to have reduced plasticity in the switch group at the second time point. Reduced plasticity of these modules was determined by the directionality of expression (up vs downregulation) of the treatment groups based on two criteria: firstly, prior to switching to the heat condition (first time point), the directionality of expression in the switch and control groups must be the same. Secondly, after switching from the heat to control condition (second time point), the directionality in the switch group must be the same as the heat group and the opposite of the control group. This would indicate that after acclimating for 24 hours, the switched expression remained in the heated state rather than reverting to pre-treatment, control state.

### Confirmation that heat treatment elicits distinct gene expression response

Heatmaps showing broad patterns of gene expression were generated using the variance stabilized counts from DESeq2. The data were plotted using the function *pheatmap::pheatmap*, clustering genes (rows) into 250 groups of similar expression using k-means algorithm.

Heatmaps were generated for time point 1 and 2 separately, though they showed little structure depending on treatment when afternoon samples were included, likely due to similarities between heat treatment and peak afternoon temperatures (Fig. S1). When separated by time point and time of day, it is visually clear that treatment was the predominant driver of gene expression at time point 2 in the morning samples (Figure 3a).

**Figure 3.**
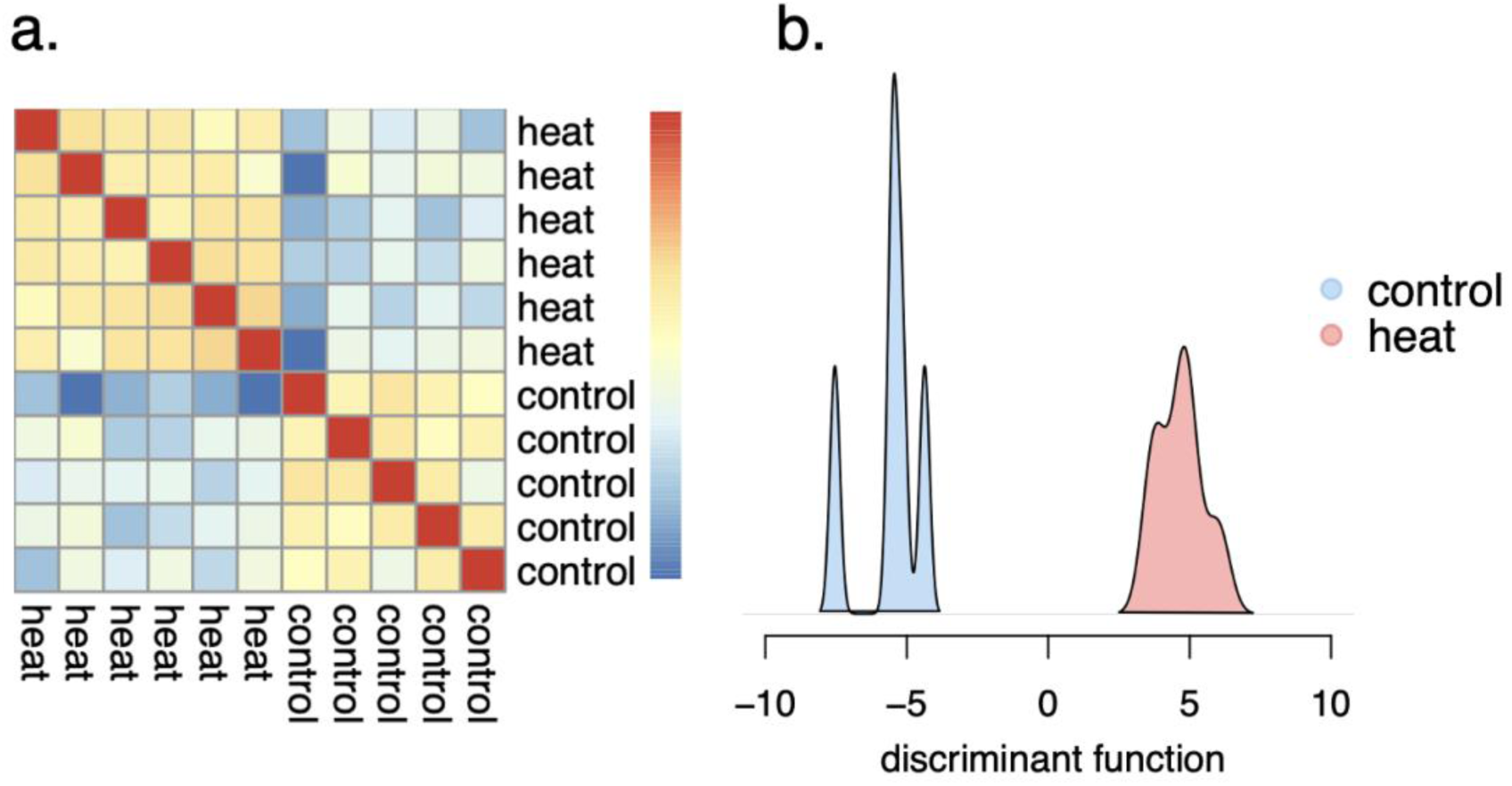
Broad patterns of gene expression in heat and control groups in the morning at time point 2. a. Heatmap showing similar (red) and different (blue) gene expression between samples. b. Colors represent heat (red) and control (blue) gene expression.

Differences between treatment groups, time points, and time of day were also investigated using discriminant analysis of principal components (DAPC). DAPC was plotted for each time point and time of day individually (Fig. S2). In accordance with the heatmaps, the difference between heat and control samples was most apparent in the morning samples at the second time point (Fig. 3b) Because of these results, only morning samples were used in treatment analyses in the main text. Analyses including afternoon samples can be found in the supplemental (Fig. S3).

## Results

### Quantifying Generalized Stress Response (GSR)

To determine if heat treatment induced a stress response, we quantified the generalized stress response represented by the “red module”, a group of 634 genes which are upregulated in response to any bleaching-level stress in *Acropora sp.* (Dixon et al., 2020). Conversely, in response to lower intensity stress, the red module is downregulated. We computed the eigengene of these genes (the first principal component of their expression variation in our samples) and found that at the second time point it was significantly downregulated in heated samples compared to controls (Fig. 4, r^2^ = 0.004), indicating that the corals were experiencing low intensity stress. Indeed, none of the fragments were visibly bleached.

**Figure 4.**
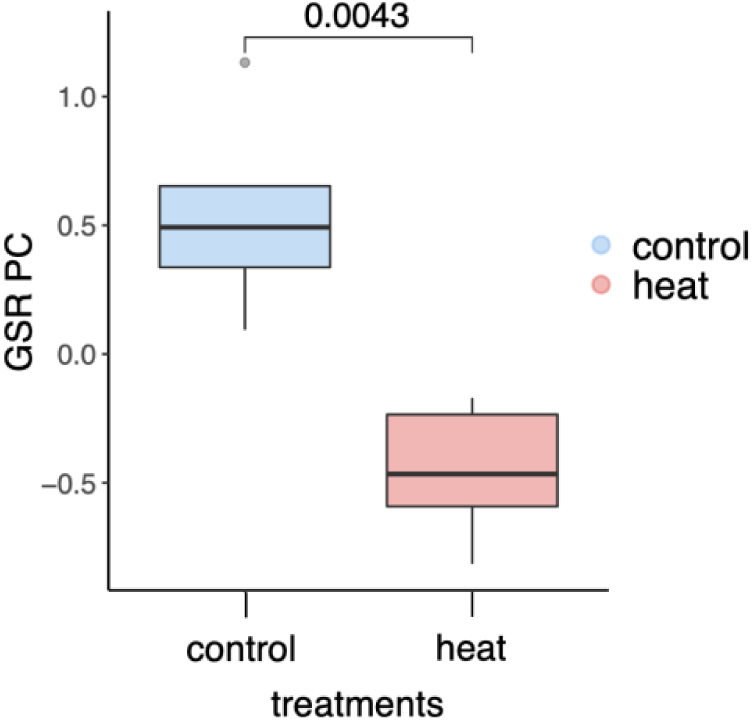
Expression of generalized stress response genes (“red module”) in control (blue) and heated (red) in the morning at the second time point.

### Baseline GBM and its association with gene expression

In invertebrates, GBM has a bimodal distribution throughout the genome (Dixon et al., 2014). Moreover, it tends to be high in genes which are constitutively and highly expressed and low in genes which are expressed at lower levels and are dynamically regulated (Sarda et al., 2012). As expected, we found the characteristic bimodal distribution of GBM levels among genes (Fig. 5a). There was a positive correlation between baseline GBM level and gene expression (r = 0.14, p = 1.99e-63; Fig. 5b), and a negative correlation between GBM and gene expression plasticity between experimental time points (r = -0.28, p = 1.77e-276; Fig. 5c).

**Figure 5.**
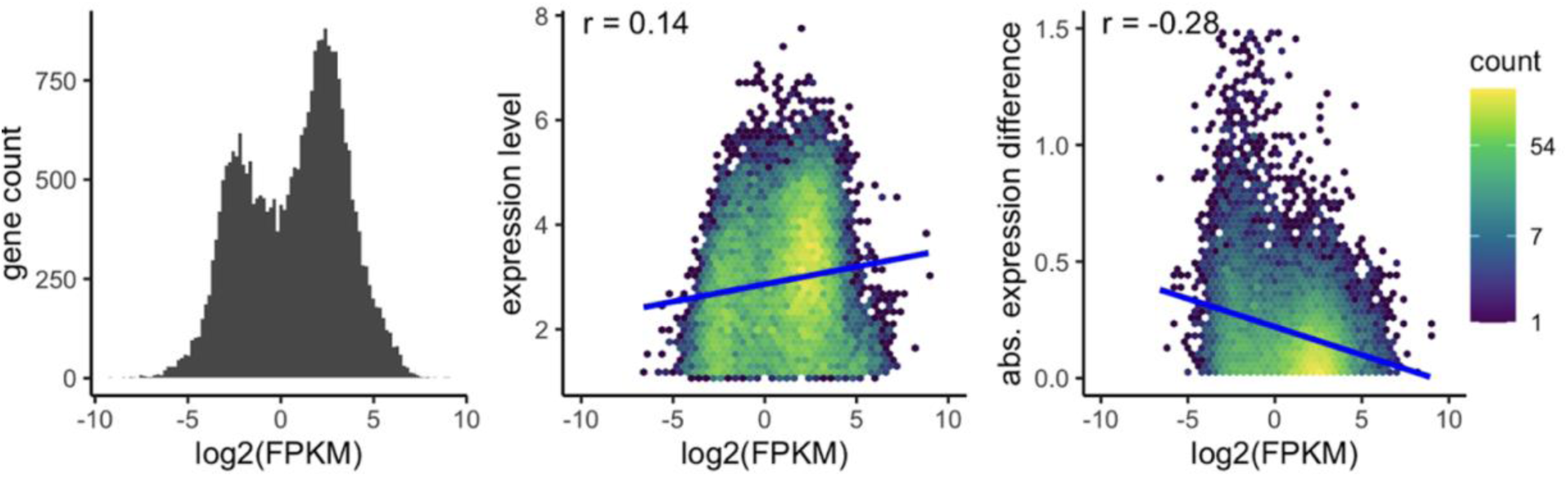
Relationship between gene expression and GBM level, quantified by reads per kilobase of exon per million reads, log_2_(FPKM). a) distribution of GBM levels across all genes. b) relationship between GBM level and gene expression level. c) relationship between GBM level and magnitude of gene expression change between experimental time points.

### Broad patterns of gene expression and GBM log-fold change

Using normalized variance-stabilized gene expression data from DESeq2, we looked for genes whose expression remained in the “heat state” 24 hours after the fragment was switched from the heat tank to the control tank (red ellipse in Fig. 6b). We then compared GBM of these genes to those whose expression reverted to the “control state” after switching (blue ellipse in Fig. 6b). Our hypothesis was that acclimatization-related GBM change would influence the gene expression plasticity, which could be a mechanism by which the organism could “learn” to maintain certain gene expression levels despite short-term environmental fluctuations.

**Figure 6.**
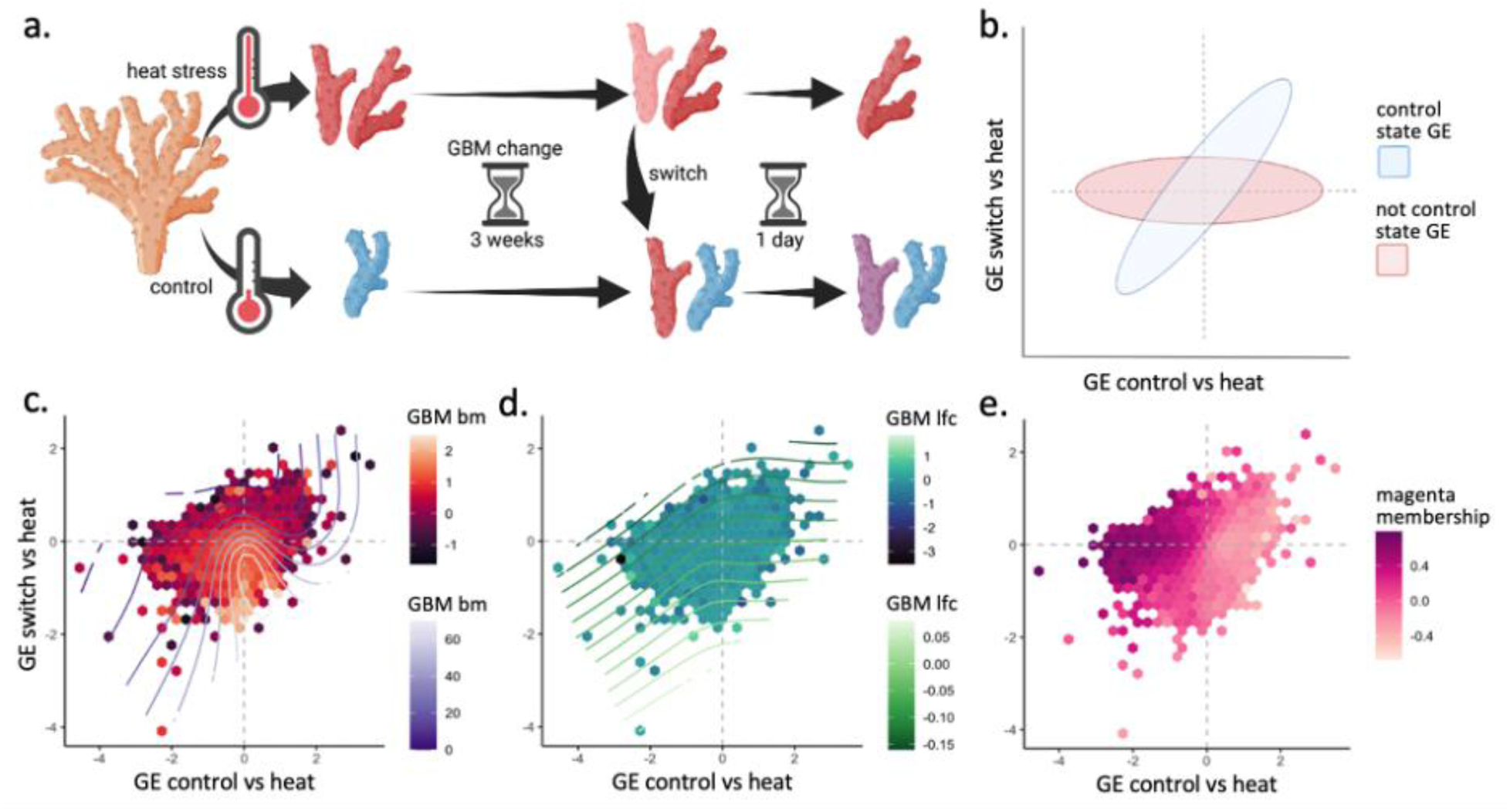
Correlation between control and switch gene expression compared to the heat group at the end of the experiment. a) Experimental design showing the control, heat, and switch groups. b) Hypothetical results showing control state gene expression (horizontal ellipse) and non-control state gene expression (diagonal ellipse). c,d,e) Experimental results showing correlation between control and switch gene expression at the end of the experiment. Hexagons represent groups of genes binned together due to similar expression. Color of the hexagons and contour lines correspond to GBM basemean (c, r^2^ = 0.058), GBM lfc (d, r^2^ = 0.002), and membership in the magenta module (e).

Overall, most of the genes in the switched group reverted to the control state (r^2^=0.135, p < 2.2e-16). However, there were also many genes that fell into the “red ellipse” region of reduced plasticity (Fig. 6b). To better visualize the relationship between GBM and plasticity, we used the ordisurf() function from the R package vegan, which fits smooth surfaces for continuous variables over ordination plots. We used GBM results from the DESeq2 model to plot contour lines representing the effect of GBM log_2_fold change and baseline GBM. Baseline GBM explained a small fraction of the variance in gene expression (Fig. 6c, r^2^ = 0.058), as expected from general association of baseline GBM with gene expression variability (Fig. 6c). Baseline GBM peaked in genes that reverted after switching but were not differentially expressed between the heat and control groups; it did not show peaks or troughs in the areas of the plot where genes with reduced gene expression plasticity were found (Fig. 6c). GBM log_2_fold change showed no association with any area of the plot and had only a negligible correlation with gene expression change (Fig. 6d, r^2^ = 0.002).

### Coexpression module with reduced plasticity

In addition to qualitatively designating genes as high or low plasticity based on their position in the scatter plot (Fig. 6b), we have applied an unsupervised gene clustering method, WGCNA (Langfelder & Horvath 2008), to group genes into coexpressed modules and looked for modules that show signatures of reduced-plasticity, i.e., differing between heat and control but not between heat and switched groups. We identified one such module, “magenta” (4252 genes, Fig. 2)

Genes with a high magenta module membership score (quantified by kME, the correlation of the gene’s expression with the module’s eigengene) formed the core of the “reduced plasticity” region of the scatter plot (Fig. 6b,e). Detecting this large group of coexpressed genes in an unsupervised fashion suggests that some common regulatory mechanism was responsible for their reduced plasticity in our experiment. However, GBM change was not significantly different than other genes in the genome (Fig. 7a).

**Figure 7.**
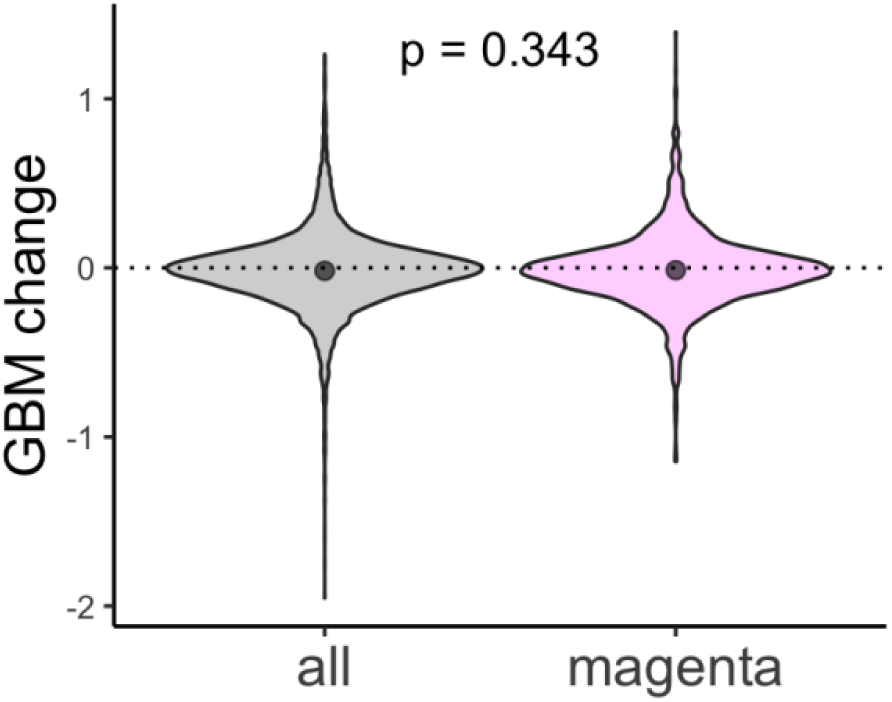
GBM change across all genes (left) and the magenta module (right). Mean GBM change in all genes (-0.017) is not significantly different from that of the magenta module (-0.0137; p=0.043).

We also considered genes which were differentially expressed in the morning versus afternoon due to daily temperature fluctuations (Fig. 8a). To measure the change in daily gene expression plasticity over the course of the experiment, we used the interaction term from our DESeq2 model, which represents the change in the magnitude of gene expression response to the afternoon temperature at the second time point vs. the first time point. . A positive correlation between this change and GBM change would indicate that plasticity increased as GBM increased (yellow and red boxes in Fig. 8b). On the other hand, a negative correlation between the two would indicate that plasticity increased as GBM increased (blue and orange boxes in Fig. 8b).

**Figure 8.**
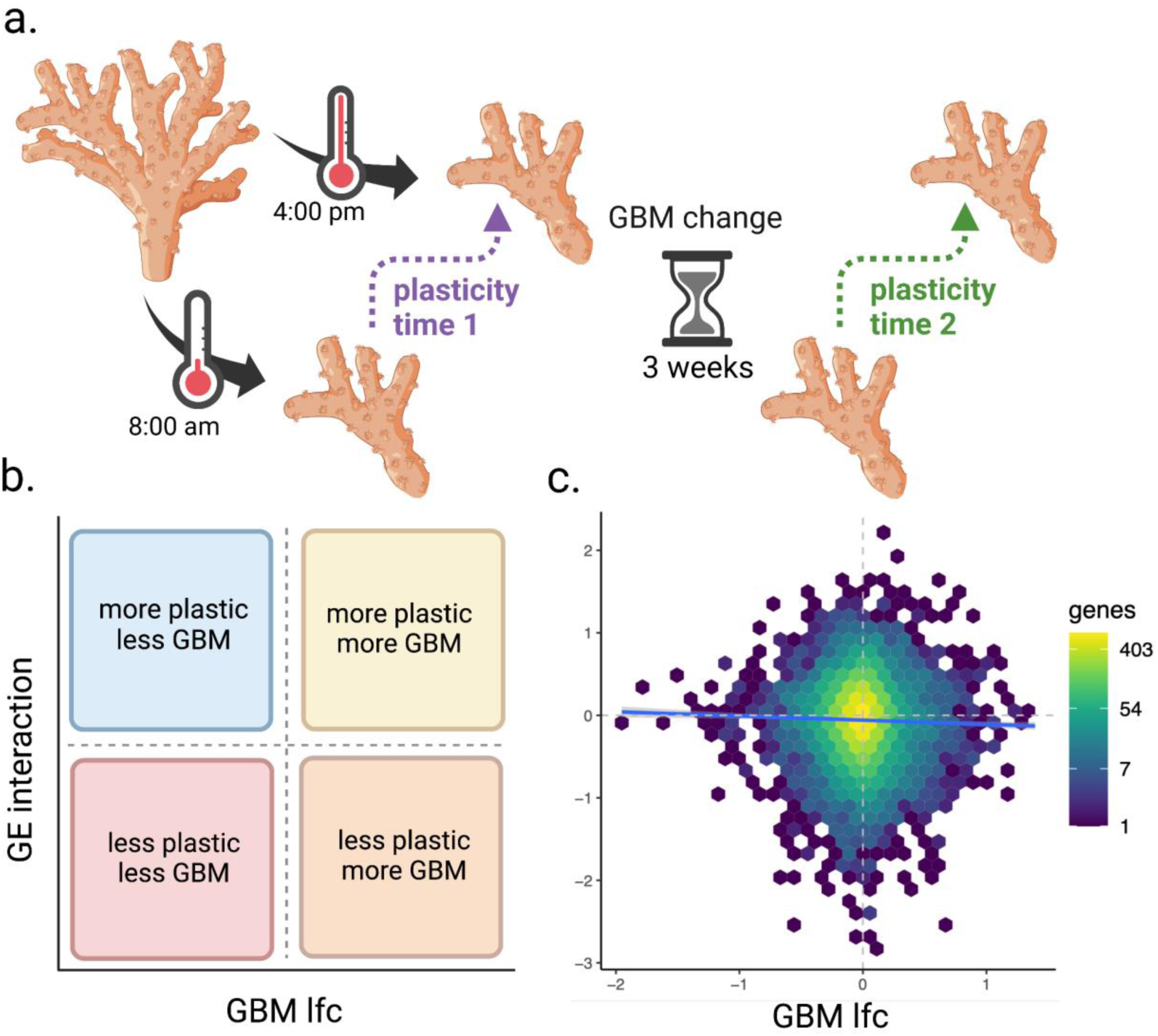
Correlation between change in daily plasticity and GBM change. a) Experimental design showing the plasticity at time 1 (purple) and time 2 (green) between 8:00am and 4:00pm. b) Hypothetical results showing the potential correlation between gene expression (effect of the interaction between experimental time point and time of day) and GBM log_2_fold change. c) Experimental results showing actual correlation between the gene expression interaction term and GBM log_2_fold change (r^2^ = 6.655e-4, p =0.004). Hexagons represent groups of genes binned together due to similar expression, and the color represents the number of genes in the hexagon.

Similar to the results from the treatment comparisons (Fig. 6), we found a negligible correlation between change in daily plasticity over time and GBM change. This relationship was very weak albeit significant, likely because of the large number of genes involved (Fig. 8; r^2^ = 6.655e-4, p =0.004).

## Discussion

Whether or not GBM plays a role in gene regulation in corals remains a matter of ongoing debate. It has been argued that gene body methylation (GBM) in invertebrates does not directly regulate gene expression since its change is neither necessary nor sufficient to induce gene expression change (Bewick et al, 2017; Harris et al., 2019; Cardoso-Júnior et al., 2021). Furthermore, the correlation between GBM change and gene expression change is typically negligible or non-existent (Dixon et al., 2022), which was also confirmed here (Fig. 6, 7). The primary objective of this work was to test whether environmentally driven changes in GBM are associated with changes in gene expression plasticity. We considered a group of genes in heat- acclimatized corals which retained heat-state expression after being returned to the control tank (the Magenta module, Fig. 6 e). Acclimatization-related GBM changes in these genes were no different from the rest of the genes in the genome (Fig. 7). We also looked at genes which were differentially regulated between the coolest and hottest time of day to see whether the magnitude of their daily plasticity change over time could be predicted by changes in GBM. We found no such association (Fig. 8 c).

If GBM does not play a role in acclimatization, then why does GBM change at all in response to the environment? It should be noted that this change is typically small compared to the change in gene expression: one study reported that even after transplanting corals across 4.5 degrees of latitude for two months, their GBM remained mostly unchanged, with more than 80% of total GBM variation attributable to differences between original coral colonies (Dixon et al., 2018). Moreover, these changes consisted in genome-wide increase or decrease of GBM disparity between the two broad gene classes, highly-methylated and lowly-methylated, rather than in targeted GBM adjustment in genes acting in specific pathways (Dixon et al., 2018). One way that environmental changes may alter broad GBM patterns is by affecting the expression and function of DNA methyltransferases. In vertebrates, changes in temperature are associated with altered expression and activity of DNMTs (Yan et al., 2015; Campos et al., 2012; Skjaerven et al., 2014). In insects it has been demonstrated that as little as a 4°C difference in temperature is correlated with changes in DNMT3 expression (Wang et al., 2020; Dai et al., 2017). If this is the case in corals, it is possible that GBM changes may simply be a consequence of spurious DNMT activity, which is consistent with our results (Fig. 6, 7).

Another possibility is that the microbiome may affect host epigenetic modifications. Corals are symbiotic organisms which harbor diverse, dynamic microbial communities (Putnam, 2021). Studies in human pathology have identified bacteria and viruses possessing proteins and enzymes, including DNMT homologs, which modulate host DNA methylation to facilitate infection (Bierne, 2017; Bierne et al., 2012; Kuss-Duerkop et al., 2018). Using metagenomes of three coral species, one study identified candidate microbes which may produce similar proteins and enzymes (Barno et al., 2021). If this is the case, it is possible that shifts in the relevant abundance of bacteria in the coral microbiome may result in GBM changes in the host. However, experimental evidence for this theory in cnidarians remains lacking.

Another pattern that remains hard to explain is the association of GBM level with broad functional gene classes: in all multicellular animals that have any GBM at all, housekeeping genes show high GBM and inducible (context-dependent) genes show low GBM (Sarda et al., 2012). This could be a simple dynamic consequence of more consistent transcription of housekeeping genes: after all, GBM is associated with transcriptional units, and it stands to reason that more consistently transcribed genome regions would attract more DNA methylation activity, likely to prevent intragenic transcription initiation (Neri et al., 2017). However, if transcription is the only GBM distribution determinant, exons should not be different from introns, yet GBM is almost always higher in exons relative to introns (Singer et al., 2015; Wang et al., 2013; Liew et al., 2018). Assuming that GBM marks are at least partially heritable (Wang et al., 2013; Liew et al., 2020), this raises an interesting possibility that GBM affects gene function and its distribution in the genome might be shaped in part by natural selection. If so, from an evolutionary standpoint GBM may be acting as a genetic mutation process, introducing heritable (and perhaps idiosyncratic) perturbations into gene function and subject to natural selection. We therefore believe that studying GBM in a population genetic /adaptation framework (rather than gene regulation /acclimatization framework) is warranted to see if GBM may be the source of adaptive, genetic variation in natural populations.

## Data availability

All scripts used in this analysis are available at https://github.com/evelynabbott/DNA_methylation_plasticity. Raw TagSeq and MdRAD reads can be accessed under the NCBI BioProject accession PRJNA1003549. The *Acropora millepora* genome is available on the Matz Lab website, https://matzlab.weebly.com/data--code.html.

## References

Barno, A. R., Villela, H. D. M., Aranda, M., Thomas, T., & Peixoto, R. S. (2021). Host under epigenetic control: A novel perspective on the interaction between microorganisms and corals. BioEssays, 43(10), Article 10. 10.1002/bies.202100068

Bewick, A. J., Sanchez, Z., Mckinney, E. C., Moore, A. J., Moore, P. J., & Schmitz, R. J. (2019). Dnmt1 is essential for egg production and embryo viability in the large milkweed bug, Oncopeltus fasciatus. Epigenetics & Chromatin, 12(1), Article 1. 10.1186/s13072-018-0246-5

Bierne, H. (2017). Cross Talk Between Bacteria and the Host Epigenetic Machinery. In W. Doerfler & J. Casadesús (Eds.), Epigenetics of Infectious Diseases (pp. 113–158). Springer International Publishing. 10.1007/978-3-319-55021-3_6

Bierne, H., Hamon, M., & Cossart, P. (2012). Epigenetics and Bacterial Infections. Cold Spring Harbor Perspectives in Medicine, 2(12), Article 12. 10.1101/cshperspect.a010272

Bird, A. (2002). DNA methylation patterns and epigenetic memory. Genes & Development, 16(1), Article 1. 10.1101/gad.947102

Campos, C., Valente, L. M. P., & Fernandes, J. M. O. (2012). Molecular evolution of zebrafish dnmt3 genes and thermal plasticity of their expression during embryonic development. Gene, 500(1), Article 1. 10.1016/j.gene.2012.03.041

Cardoso-Júnior, C. A. M., Yagound, B., Ronai, I., Remnant, E. J., Hartfelder, K., & Oldroyd, B. P. (2021). DNA methylation is not a driver of gene expression reprogramming in young honey bee workers. Molecular Ecology, 30(19), Article 19. 10.1111/mec.16098

Chandler, V. L., & Walbot, V. (1986). DNA modification of a maize transposable element correlates with loss of activity. Proceedings of the National Academy of Sciences, 83(6), Article 6. 10.1073/pnas.83.6.1767

Dai, T.-M., Lü, Z.-C., Wang, Y.-S., Liu, W.-X., Hong, X.-Y., & Wan, F.-H. (2018). Molecular characterizations of DNA methyltransferase 3 and its roles in temperature tolerance in the whitefly, Bemisia tabaci Mediterranean. Insect Molecular Biology, 27(1), Article 1. 10.1111/imb.12354

de Mendoza, A., Lister, R., & Bogdanovic, O. (2020). Evolution of DNA Methylome Diversity in Eukaryotes. Journal of Molecular Biology, 432(6), Article 6. 10.1016/j.jmb.2019.11.003

Dixon, G., Abbott, E., & Matz, M. (2020). Meta-analysis of the coral environmental stress response: Acropora corals show opposing responses depending on stress intensity. Molecular Ecology, 29(15), Article 15. 10.1111/mec.15535

Dixon, G. B., Bay, L. K., & Matz, M. V. (2014). Bimodal signatures of germline methylation are linked with gene expression plasticity in the coral Acropora millepora. BMC Genomics, 15(1), Article 1. 10.1186/1471-2164-15-1109

Dixon, G., Liao, Y., Bay, L. K., & Matz, M. V. (2018). Role of gene body methylation in acclimatization and adaptation in a basal metazoan. Proceedings of the National Academy of Sciences, 115(52), Article 52. 10.1073/pnas.1813749115

Dixon, G., & Matz, M. (2022). Changes in gene body methylation do not correlate with changes in gene expression in Anthozoa or Hexapoda. BMC Genomics, 23(1), Article 1. 10.1186/s12864-022-08474-z

Epigenome-associated phenotypic acclimatization to ocean acidification in a reef-building coral | Science Advances. (2023, March 8). https://www.science.org/doi/full/10.1126/sciadv.aar8028

Fuller, Z. L., Mocellin, V. J. L., Morris, L., Cantin, N., Shepherd, J., Sarre, L., Peng, J., Liao, Y., Pickrell, J., Andolfatto, P., Matz, M., Bay, L. K., & Przeworski, M. (2019). Population genetics of the coral Acropora millepora: Towards a genomic predictor of bleaching. bioRxiv. 10.1101/867754

Gatzmann, F., Falckenhayn, C., Gutekunst, J., Hanna, K., Raddatz, G., Carneiro, V. C., & Lyko, F. (2018). The methylome of the marbled crayfish links gene body methylation to stable expression of poorly accessible genes. Epigenetics & Chromatin, 11(1), Article 1. 10.1186/s13072-018-0229-6

Genome-Wide Evolutionary Analysis of Eukaryotic DNA Methylation | Science. (2023, March 8). https://www.science.org/doi/10.1126/science.1186366

Hackett, J. A., & Surani, M. A. (2013). DNA methylation dynamics during the mammalian life cycle. Philosophical Transactions of the Royal Society B: Biological Sciences, 368(1609), Article 1609. 10.1098/rstb.2011.0328

Harris, K. D., Lloyd, J. P. B., Domb, K., Zilberman, D., & Zemach, A. (2019). DNA methylation is maintained with high fidelity in the honey bee germline and exhibits global non-functional fluctuations during somatic development. Epigenetics & Chromatin, 12(1), Article 1. 10.1186/s13072-019-0307-4

J, O. (2010). vegan: Community Ecology Package. http://CRAN.R-Project.Org/Package=vegan. https://cir.nii.ac.jp/crid/1571980076147214336

Jones, P. A. (2012a). Functions of DNA methylation: Islands, start sites, gene bodies and beyond. Nature Reviews Genetics, 13(7), Article 7. 10.1038/nrg3230

Jones, P. A. (2012b). Functions of DNA methylation: Islands, start sites, gene bodies and beyond. Nature Reviews Genetics, 13(7), Article 7. 10.1038/nrg3230

Kuss-Duerkop, S. K., Westrich, J. A., & Pyeon, D. (2018). DNA Tumor Virus Regulation of Host DNA Methylation and Its Implications for Immune Evasion and Oncogenesis. Viruses, 10(2), Article 2. 10.3390/v10020082

Li, H., Handsaker, B., Wysoker, A., Fennell, T., Ruan, J., Homer, N., Marth, G., Abecasis, G., Durbin, R., & 1000 Genome Project Data Processing Subgroup. (2009). The Sequence Alignment/Map format and SAMtools. Bioinformatics, 25(16), Article 16. 10.1093/bioinformatics/btp352

Liao, Y., Smyth, G. K., & Shi, W. (2014). featureCounts: An efficient general purpose program for assigning sequence reads to genomic features. Bioinformatics, 30(7), Article 7. 10.1093/bioinformatics/btt656

Liew, Y. J., Howells, E. J., Wang, X., Michell, C. T., Burt, J. A., Idaghdour, Y., & Aranda, M. (2020). Intergenerational epigenetic inheritance in reef-building corals. Nature Climate Change, 10(3), Article 3. 10.1038/s41558-019-0687-2

Liew, Y. J., Zoccola, D., Li, Y., Tambutté, E., Venn, A. A., Michell, C. T., Cui, G., Deutekom, E. S., Kaandorp, J. A., Voolstra, C. R., Forêt, S., Allemand, D., Tambutté, S., & Aranda, M. (2018). Epigenome-associated phenotypic acclimatization to ocean acidification in a reef-building coral. Science Advances, 4(6), Article 6. 10.1126/sciadv.aar8028

Love, M. I., Huber, W., & Anders, S. (2014). Moderated estimation of fold change and dispersion for RNA-seq data with DESeq2. bioRxiv. 10.1101/002832

Lyko, F. (2018). The DNA methyltransferase family: A versatile toolkit for epigenetic regulation. Nature Reviews Genetics, 19(2), Article 2. 10.1038/nrg.2017.80

Martin, M. (2011). Cutadapt removes adapter sequences from high-throughput sequencing reads. EMBnet.Journal, 17(1), Article 1. 10.14806/ej.17.1.200

Moore, L. D., Le, T., & Fan, G. (2013). DNA Methylation and Its Basic Function. Neuropsychopharmacology, 38(1), Article 1. 10.1038/npp.2012.112

Neri, F., Rapelli, S., Krepelova, A., Incarnato, D., Parlato, C., Basile, G., Maldotti, M., Anselmi, F., & Oliviero, S. (2017). Intragenic DNA methylation prevents spurious transcription initiation. Nature, 543(7643), Article 7643. 10.1038/nature21373

Putnam, H. M. (2021). Avenues of reef-building coral acclimatization in response to rapid environmental change. Journal of Experimental Biology, 224(Suppl_1), Article Suppl_1. 10.1242/jeb.239319

Sarda, S., Zeng, J., Hunt, B. G., & Yi, S. V. (2012). The Evolution of Invertebrate Gene Body Methylation. Molecular Biology and Evolution, 29(8), Article 8. 10.1093/molbev/mss062

Singer, M., Kosti, I., Pachter, L., & Mandel-Gutfreund, Y. (2015). A diverse epigenetic landscape at human exons with implication for expression. Nucleic Acids Research, 43(7), Article 7. 10.1093/nar/gkv153

Smith, Z. D., & Meissner, A. (2013). DNA methylation: Roles in mammalian development. Nature Reviews Genetics, 14(3), Article 3. 10.1038/nrg3354

Wang, Y., Wang, F., Hong, D., Gao, S., Wang, R., & Wang, J. (2020). Molecular characterization of DNA methyltransferase 1 and its role in temperature change of armyworm Mythimna separata Walker. Archives of Insect Biochemistry and Physiology, 103(4), Article 4. 10.1002/arch.21651

Yan, X., Liu, H., Liu, J., Zhang, R., Wang, G., Li, Q., Wang, D., Li, L., & Wang, J. (2015). Evidence in duck for supporting alteration of incubation temperature may have influence on methylation of genomic DNA. Poultry Science, 94(10), Article 10. 10.3382/ps/pev201

