## Supplementary material for "Modifications to gene body methylation does not alter gene expression plasticity in a reef-building coral": (Fig. S3).

Figure S3.

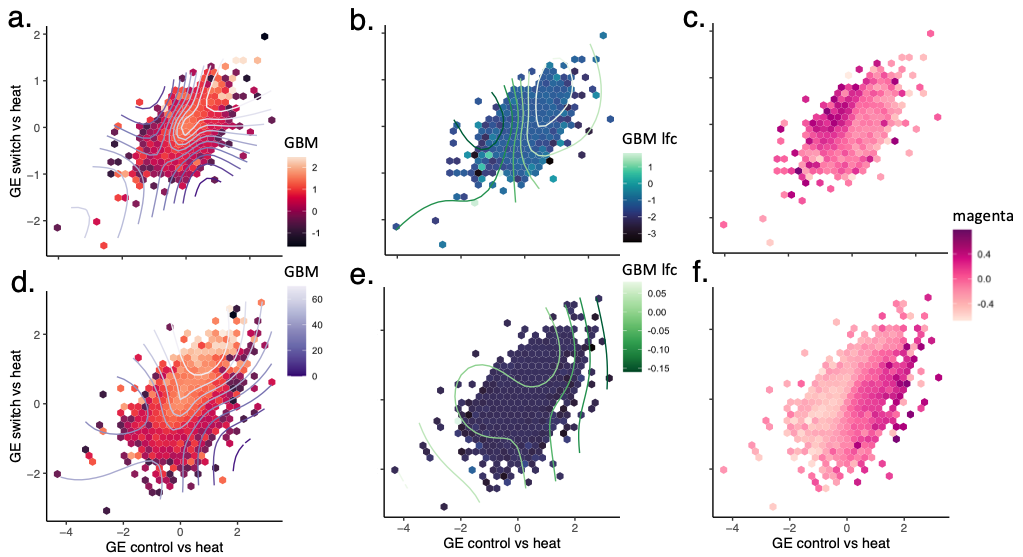

**Figure S3.** Experimental results showing correlation between control and switch gene expression at the end of the experiment, including afternoon samples (a-c) and excluding morning samples (d-f). Hexagons represent groups of genes binned together due to similar expression. a-c) Including all samples, the color of the hexagons and contour lines correspond to GBM basemean (a, r^2^ = 0.041, p <2e-16), GBM lfc (b, r^2^ = -5.61e-08, p = 0.719), and membership in the magenta module (c). d-f) Including only afternoon samples, the color of the hexagons and contour lines correspond to GBM basemean (d, r^2^ = 0.084, p <2e-16), GBM lfc (e, r^2^ = 0.003, p = 7.59e-06), and membership in the magenta module (f).
