## Supplementary material for "Modifications to gene body methylation does not alter gene expression plasticity in a reef-building coral": (Fig. S2)

Figure S2.

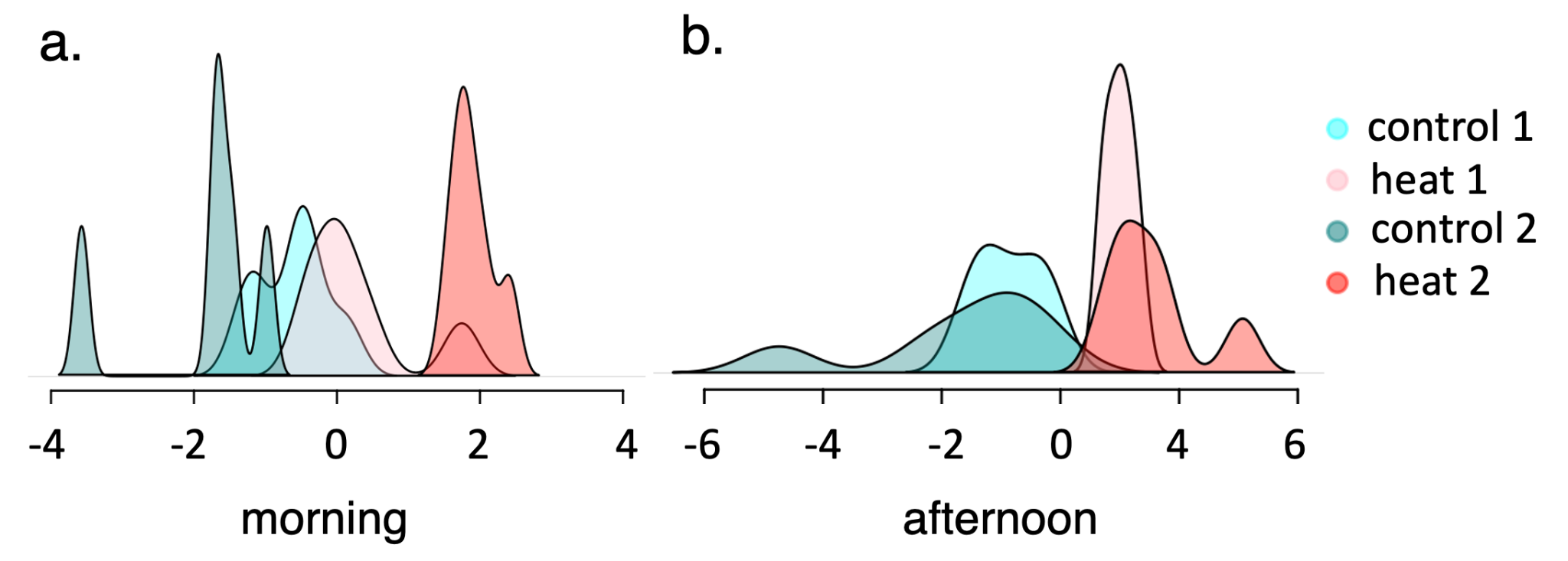

Figure S2. Discriminant analysis of principal components (DAPC) plotted for each time point in the morning (a) and afternoon (b). Colors represent control treatments (light teal for time point 1, dark teal for time point 1) and heat treatments (pink for time point 1, red for time point 2).
