## Supplementary material for "Modifications to gene body methylation does not alter gene expression plasticity in a reef-building coral": (Fig. S1)

Figure S1.

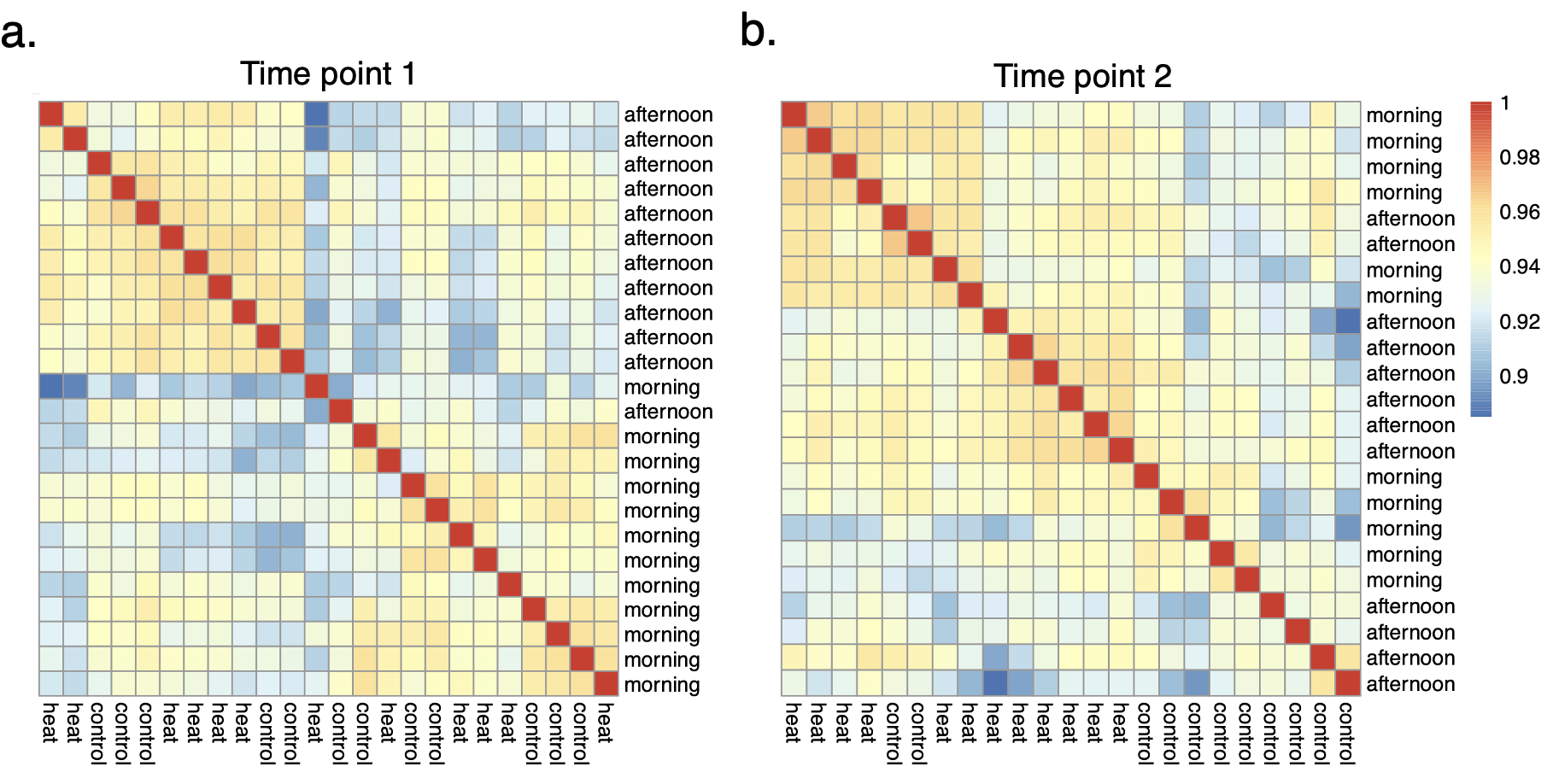

Figure S1. Broad patterns of gene expression in heat and control groups in the morning and afternoon at time point (a) and time point 2 (b). Heatmaps show similar (red) and different (blue) gene expression between samples.
